# The effect of variable inertial resistance on force, velocity, power and muscle activation during a chest press exercise

**DOI:** 10.1101/2023.12.16.572002

**Authors:** Luca Zoffoli, Andrea Biscarini, Silvano Zanuso

## Abstract

This study investigated the effect of the inertial component of the resistance (*INERTIA*) at different intensity levels (*LOAD* ) on force (*FORCE* ), velocity (*VELOCITY* ), power (*POWER*), and the muscle activity of the pectoralis major (*EMG*_*PM*_), anterior deltoid (*EMG*_*DA*_) and triceps brachii (*EMG*_*TB*_) muscles during a chest press exercise.

A motor-driven exercise apparatus was programmed to offer resistance with different inertial profiles over the range of movement (*ROM* ): gravitational-type constant inertia (*I*_*FULL*_); no-inertia (*I*_*ZERO*_); linearly descending inertia along the *ROM* (*I*_*VAR*_). Nine healthy adults performed five, maximal-effort, explosive movements with each inertial profile at 30, 50 and 70% of their 1 repetition maximum. Meanwhile, the *EMG*_*PM*_, *EMG*_*DA*_ and *EMG*_*TB*_ signals were obtained jointly with the *FORCE, VELOCITY* and *POWER* readings returned by the exercise apparatus.

One-dimensional statistical non-parametric maps based on 2-way repeated measures ANOVA (*SnPM* ) were employed to evaluate the effect of *LOAD* and *INERTIA* on the collected timeseries. Paired t-tests were then used as post-hoc tests on the portions of the *ROM* denoting significant differences in the *SnPM*.

Higher *LOAD* resulted in elevated outcomes over large portions of the *ROM* in all the investigated timeseries. Compared to *I*_*FULL*_, *I*_*ZERO*_ allowed greater *VELOCITY* at the cost of lower *FORCE* throughout the *ROM*, while *I*_*VAR*_, despite the lower *VELOCITY* than *I*_*ZERO*_, resulted in higher *FORCE* and *POWER* output. In addition, *I*_*ZERO*_ and *I*_*VAR*_ elevated *EMG*_*TB*_ at the end of the *ROM* with respect to *I*_*FULL*_. *I*_*VAR*_ overcame both *I*_*FULL*_ and *I*_*ZERO*_ in terms of *FORCE* and *POWER*, which indicates that variable inertial profiles might be effectively integrated into resistance exercise programs. Ultimately, this study suggested that *INERTIA* acts independently to the imposed *LOAD* on the *FORCE, VELOCITY* and *POWER* production. Coaches and therapists are encouraged to account for the type of *INERTIA* as one of the parameters considered during the exercise selection for their athletes or patients.

## Introduction

Resistance training includes any physical activity performed against an opposing force (resistance). Despite the several types of resistances investigated [1], the commercial availability of free-weights, weight-stack and plate-loaded [2] machines allowed the large application of gravitational and isotonic-like resistances in sport [3] and fitness [4], with a huge potential also for rehabilitation purposes [5]. In general, free-weights are thought to provide constant (gravitational) resistance to the movement, while weight stack machines, thanks to the presence of specifically designed asymmetric pulleys, are deemed to generate isotonic resistance profiles. However, both concepts can be misleading as they assume that the movement occurs at constant velocity, thus neglecting the impact of the inertia to the overall resistance. Inertial force strictly depends on the amplitude of the acceleration imposed on the lifted mass [6], and therefore becomes particularly relevant when different movements speeds are compared.

Lifting weights at high velocity is usually recommended for the development of strength and power [8]. According to Newton’s laws [7], this entails high accelerations at the beginning of the movement, which translates to a relevant amount of inertial force that must be won to effectively achieve high velocities. Conversely, during the later phase of the movement, the inertial force lowers the total resistance opposed to the movement by an amount proportional to the magnitude of the deceleration necessary to halt the weight at the end of the lift. This behaviour has important consequences on both acute and chronic training adaptations. For instance, either fast or moderate-slow resistance training routines induce comparable muscular strength gains [9]. However, moving at higher velocity increases the number of performed repetitions before failure [10] and elevates the surface electromyographic activity (*EMG* ) of the *pectoralis major* muscle [11].

Some authors suggested that non-inertial resistances can be effective for strength and power development. When gravitational and pneumatic resistance have been compared, the latter resulted in higher movement velocities and muscle activation [12]. Likewise, elastic resistance has shown significant improvements in strength and power gains over standard free-weight exercises [13]. Broadly speaking, the absence of inertia allows the achievement of higher movement velocity at the cost of reduced force production [12]. However, theoretical models suggested that this behaviour might not be optimal to maximize the force production and rate of force development, which could be improved through the application of variable inertia profiles [14].

So far, the study of different inertial profiles during resistance exercise has been limited to the comparison of constant and non-inertial resistances. In addition, these studies have been often impaired by the use of different exercise apparatuses to elicit the target inertial resistances. This produces a challenge when needing to isolate the inertial effects from those derived by the specific mechanics of the exercise equipment. Consequently, this study employed the use of a novel exercise equipment capable of providing programmable resistive profiles, thus allowing to precisely isolate the inertial effects to those deriving from the movement mechanics or the overall exercise intensity. This was used to introduce a novel variable resistance profile to compare with more conventional constant and non-inertial resistances on force, velocity, power and muscle activation of the prime movers during a conventional chest press exercise.

## Materials and methods

### Subjects

Nine healthy males (Age: 41.8 *±* 10.7 *years*; Weight: 83.8 *±* 5.8 *kg*; 1 Repetition Maximum: 98.4 *±* 23.3 *kg*) with advanced experience in resistance training were enrolled, on voluntary basis from the 3^rd^ to the 30^th^ of June 2022. All participants were subject to a medical screening for the practice of non-competitive physical exercise within the 6 months before the start of the study. The participants were extensively briefed about the study modalities, purpose, and experimental hypothesis before starting the tests. The study was approved by the Ethical committee of the University of Perugia (protocol number 130298) and written informed consent was obtained from each participant before the test. Eligibility criteria for this study were: more than 2 years of regular resistance training experience; absence of functional or injury related limitations to the movement studied (chest press).

### Exercise apparatus

A commercially available electro-mechanically driven exercise apparatus (Biostrength™ Chest press, Technogym SPA, Cesena, Italy) was used (Fig. 1). This type of exercise equipment has been already employed by research studies [15] and was chosen because it allowed the design of arbitrary resistance profiles. Therfore making possible to generate resistance with variable levels of inertia within the range of movement (*ROM* ). In addition, this equipment could record the force and displacement generated during the exercise at sampling rate of 20 Hz with 16-bit resolution.

**Fig 1.**
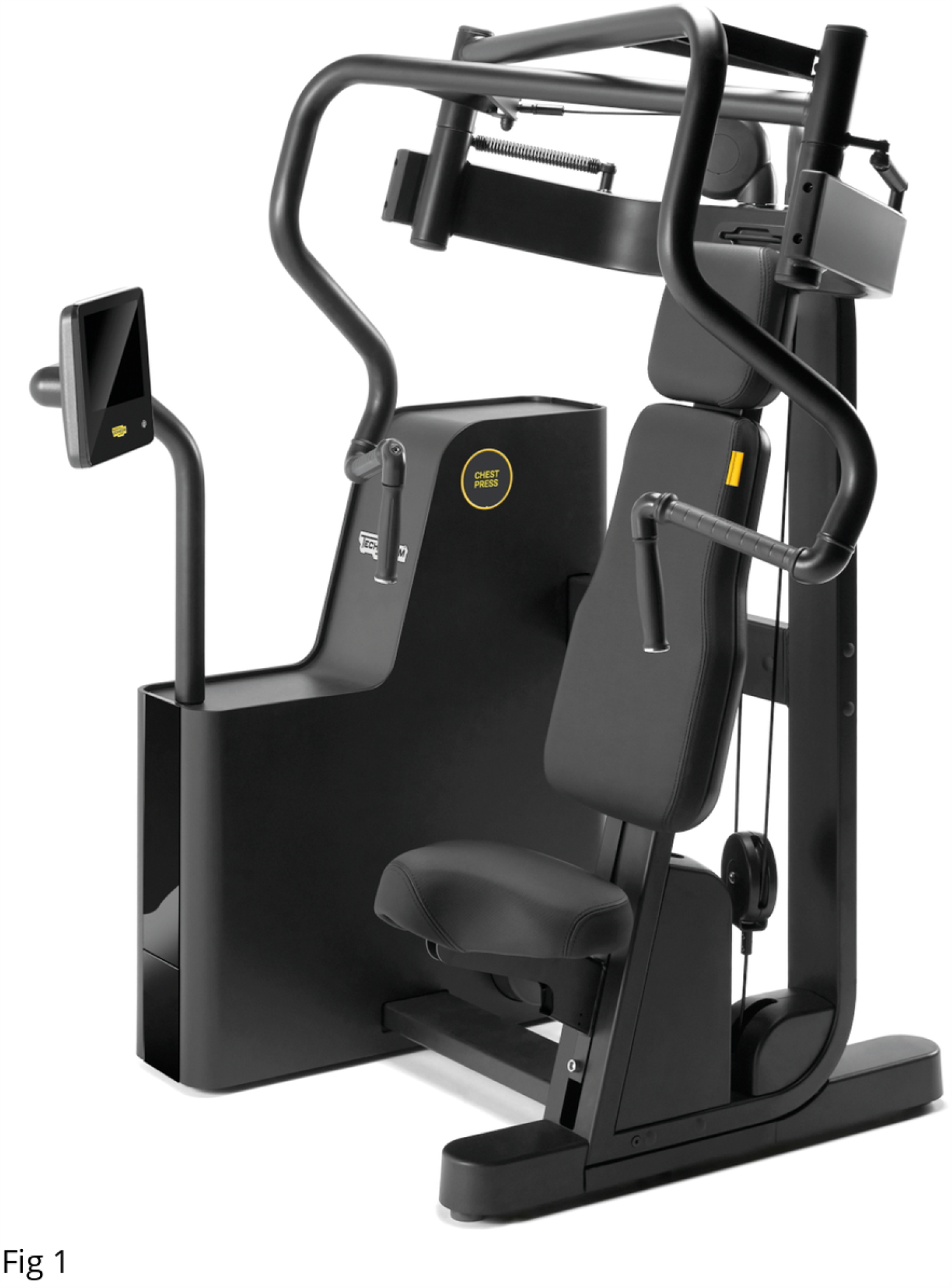
Exercise apparatus. Technogym Biostrength™ Chest Press

## Experimental setup

Three distinct inertial resistance profiles have been designed to reflect specific physical behaviours (see the S1 Appendix for a detailed mathematical description):

- *I*_*FULL*_ provided the inertial profile corresponding to the use of a conventional weight stack as resistance.
- *I*_*ZERO*_ denoted the complete absence of inertial resistance. Only the selected weight component was effectively offered during the exercise.
- *I*_*VAR*_ represented a variable inertial profile linearly reducing from *I*_*FULL*_ to *I*_*ZERO*_ throughout the 25-75% range of the *ROM*.

Pairs of unipolar, Ag/AgCl, ∅ = 24 *mm*, pre-gelled, disposable *EMG* electrodes (Covidien Kendall, Minneapolis, MN, USA) were secured on the right side of the body over the *pectoralis major, anterior deltoid* and the *long head of the triceps brachii* muscles [16]. These muscles were chosen as they are prime movers for the studied exercise [17]. The *EMG* electrodes were placed parallel to the muscle fibres with 2 *cm* inter-electrode distance. Before their placement, the skin was shaved, slightly abraded via sandpaper, and cleaned with an alcoholic solution. All the *EMG* data (input impedance: 100 *M Ω*; CMRR: *>>* 110 *dB*; baseline noise: *<* 1 *µV* ; gain: 240.6) were synchronously sampled at 1 *kHz* and stored on a computer using a 16 *bit* resolution wireless system (FreeEMG, BTS Bioengineering SPA, Italy).

Pairs of semi-spherical reflective markers (∅ = 10 *mm*) were secured to the ends of the exercise apparatus handles. Kinematic data were collected via a 10 cameras motion capture system (SMART DX-7000, BTS Bioengineering SPA, Italy), time synchronised with the *EMG* sensors. The kinematic data were sampled at 500 *Hz* with 4 *Mpixel* resolution. Calibration of the kinematic system resulted in a measurement error*<* 0.3 *mm* for all cameras.

## Testing procedure

The participants performed a general warm-up of 5 minutes on a cycle ergometer (Skillbike™, Technogym SPA, Cesena, Italy) at self-selected resistance. Afterwards, according to the guidelines of the exercise apparatus manufacturer, the participants sat on the exercise apparatus and regulated the seat height to have the centre of their shoulder joints at the height of a pair of yellow flags secured on the back rest. This ensured that the participants performed the exercise with the upper limbs almost aligned with the rotation plane of the handles, allowing the optimal generation of force. Three repetitions were performed at the minimal load (10 *kg*) at slow constant pace. Approximately 3 seconds were allowed for the concentric and eccentric phase, with no isometric rest in-between.

After the warm-up, the participants were asked to perform 6 repetitions with self-selected load using the *I*_*FULL*_ resistance profile. After 3 minutes of rest, the load was increased, and another 6 repetitions were performed. This procedure was repeated, usually 2-3 times, until the participant was no longer able to perform the six repetitions. Then, the participant’s 1 Repetition Maximum (*1RM* ) was estimated according to the load and the number of repetitions performed during the last set [18]. After another 3 minutes of rest, three maximal isokinetic, concentric-only repetitions were performed. The movement speed was set at approximately 0.24 *m/s* (measured at the handles mid-points) to mimic the average lifting speed of the *1RM* in the bench press exercise [19].

Next, the participants performed 9 sets of 5 repetitions at maximal concentric speed, each with a specific combination of the three inertial profiles (*I*_*FULL*_, *I*_*ZERO*_, *I*_*VAR*_) and three intensity levels: 30%, 50% and 70% of their *1RM* (namely, *1RM*_*30*_, *1RM*_*50*_ and *1RM*_*70*_). Every set was separated by a minimum of 3 minutes of recovery, and the adopted inertial condition was randomized for each participant. During the tests, the participants were instructed to *”push as much and as fast as possible”*, and they were not aware of the specific inertial profile being used.

## Data processing

Each *EMG* signal collected from the right *pectoralis major* (*EMG*_*PM*_), *anterior deltoid* (*EMG*_*DA*_) and *triceps brachii* (*EMG*_*TB*_) muscles were centered around their mean and filtered by a fourth order, band-pass, zero-phase, Butterworth filter with bandwidth of 20 Hz [20] and 450 Hz [16] before being full-wave rectified. For each trial, the force generated by the user over time (*FORCE*)was obtained from the motor readings. Conversely, the instantaneous velocity (*VELOCITY* ) was obtained derivating the right handle mid-point position over time [21]. All data were then linear enveloped by means of a fourth order, low-pass, phase-corrected, Butterworth filter with cut-off frequency of 3 Hz [22].

*FORCE* data were time synchronized with *VELOCITY* and *EMG* data detecting the time lag corresponding to the peak in the cross-correlation between the positional readings of the exercise apparatus with those of the right handle mid-point obtained from kinematic acquisitions. Ultimately, the power output (*POWER*) was calculated as the product between *FORCE* and *VELOCITY*.

The time instants corresponding to the beginning and end of the concentric phase of each repetition were obtained by:

1. looking at the local minima and maxima in the anterior-posterior displacement of the handles;
2. identifying the closest zero-crossing sample in the *VELOCITY* signal to each detected minima and maxima.

For each trial, the first and last repetition were excluded from the analysis to avoid possible confounding effects due to the initiation or end of the movement. Then, for each investigated muscle, the processed *EMG* signals were normalized by the mean of the peak values resulting from the repetitions performed with isokinetic resistance.

Finally, for each detected concentric phase, the *EMG*_*PM*_, *EMG*_*DA*_, *EMG*_*TB*_, *FORCE, VELOCITY* and *POWER* data have been cubic-spline interpolated to 101 samples.

## Statistical analysis

Because of the one-dimensional nature of the investigated data, to evaluate the effects of the lifted load (*LOAD* ) and the inertial profile (*INERTIA*) on each investigated parameter, statistical maps based on 2 way repeated measures ANOVA were generated [23]. During exploratory analysis, Bayesian Information Criterion [24] maps were obtained for every parameter through the calculation of this score for each of the 101 samples defining the signals. The median value of each map was then calculated. Lower median values were obtained from the models excluding the interaction term, therefore each parameter has been analysed considering only the main effects. Similar maps were obtained applying the Shapiro-Wilk test, which denoted the lack of normality on several portions of the data. Hence, a non parametric approach based on permutation tests was used to detect the regions of the *ROM* highlighted by significant differences [25]. *η*^2^ maps were also calculated. When significant differences in specific regions of the *ROM* were found, the *ROM* _START:STOP_ notation has been used to provide a concise representation of the significant regions emerged from the statistical analysis. For each significant region of the *ROM*, the mean *η*^2^ (namely 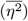) was calculated as measure of effect size. Posthoc analyses were then performed via permutation tests based on paired t-tests between the mean value obtained from the groups being compared over the significant region. For each posthoc comparison, the mean values with 95% confidence intervals of the compared quantities was reported.

The level of significance (*α*) was set at 0.05 and 10000 permutations were used to derive the distrubution from which the *p* values have been extracted. The Holm-Sidak correction [26] was employed to correct for the multiple comparisons during posthoc analyses.

All data were reduced, processed and analysed via custom python scripts (version 3.11.3, https://www.python.org) with the use of the libraries *numpy* (version 1.25, https://www.numpy.org), *matplotlib* (version 3.7.0, https://www.matplotlib.org), *pandas* (version 2.0.3, https://pandas.pydata.org), *scipy* (version 1.11.1,https://www.scipy.org), *spm1d* (version 0.4.18, https://www.spm1d.org) and *statsmodels* (version 0.14.0, https://www.statsmodels.org).

## Results

*FORCE* was increased by *LOAD* over the whole *ROM* (Fig. 2), with higher values corresponding to higher *1RM* percentages. Conversely, *INERTIA* altered the *FORCE* production during the mid-range of the concentric phase (*ROM* _18%:79%_), with the posthoc analysis revealing progressively higher values for *I*_*ZERO*_, *I*_*FULL*_ and *I*_*VAR*_, respectively (Table1).

**Fig 2.**
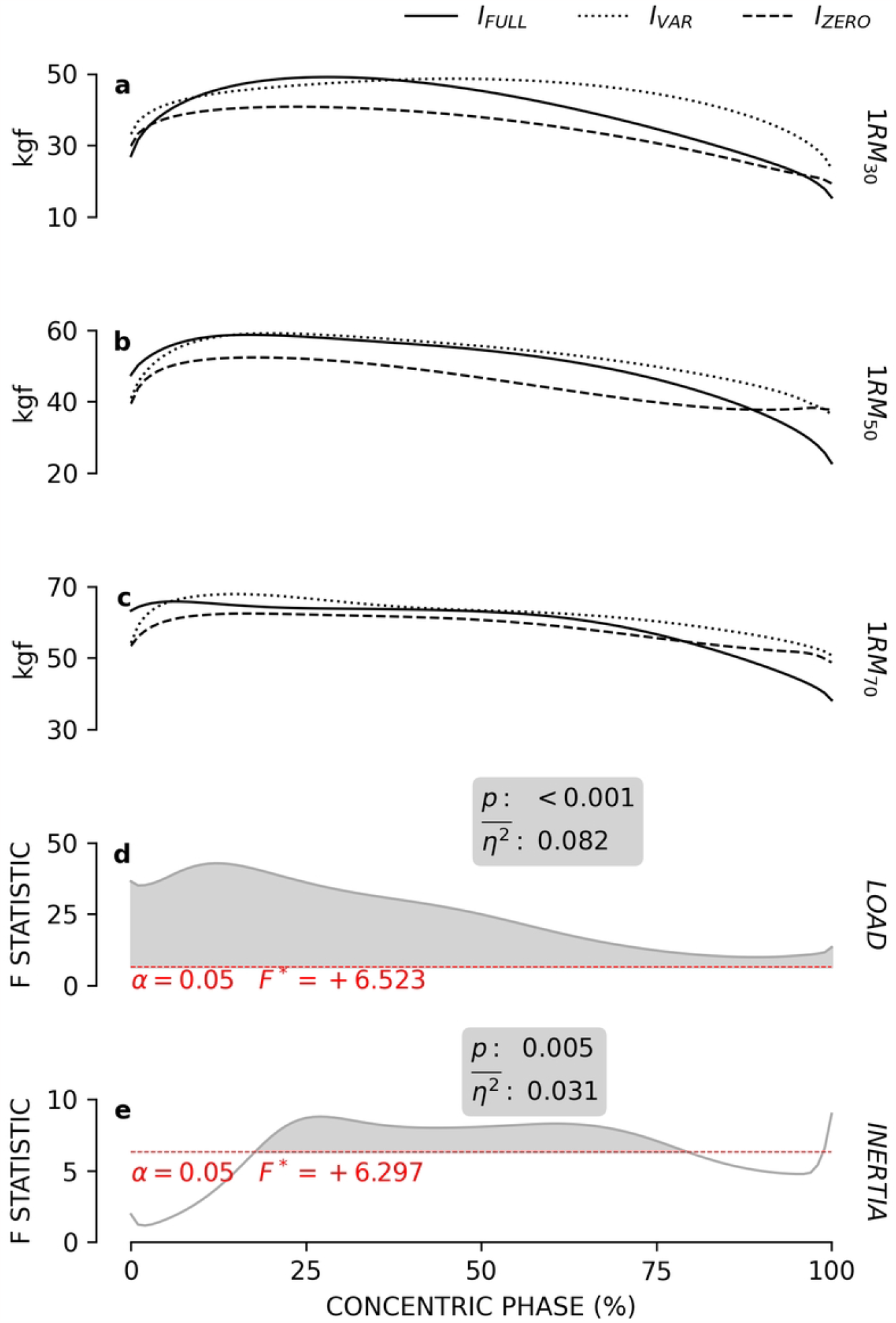
*FORCE* patterns. The subfigures **a, b** and **c** respectively report the mean force patterns obtained at 30, 50 and 70% of the *1RM* for each inertial profile. The grey lines in subfigures **d** and **e** show the statistical maps for the *LOAD* and *INERTIA* effects. The red dashes indicate the critical F value corresponding to *α*. The grey areas highlight the significant regions of the map and the boxes report the calculated *p* values and effect size 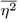.

*VELOCITY* progressively decreased over almost all the concentric phase (*ROM* _0%:96%_) as *LOAD* increased (Fig. 3). Also *INERTIA* had a significant effect on the major part of the concentric phase (*ROM* _3%:100%_), with progressively reduced *VELOCITY* values respectively for *I*_*ZERO*_, *I*_*VAR*_ and *I*_*FULL*_ (Table 1).

**Table 1.**
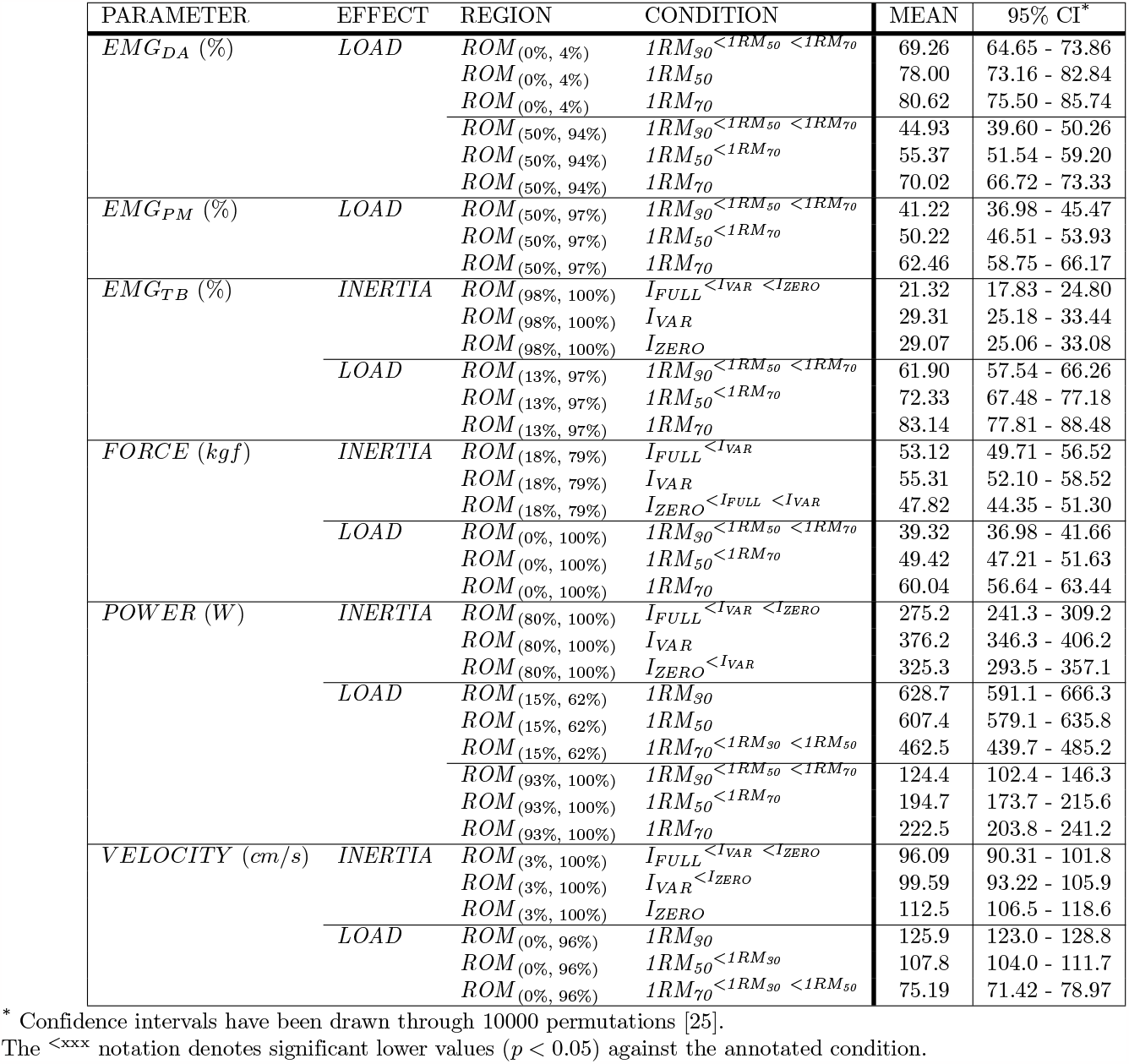
LOAD and INERTIA comparisons over significant regions of the ROM.

**Fig 3.**
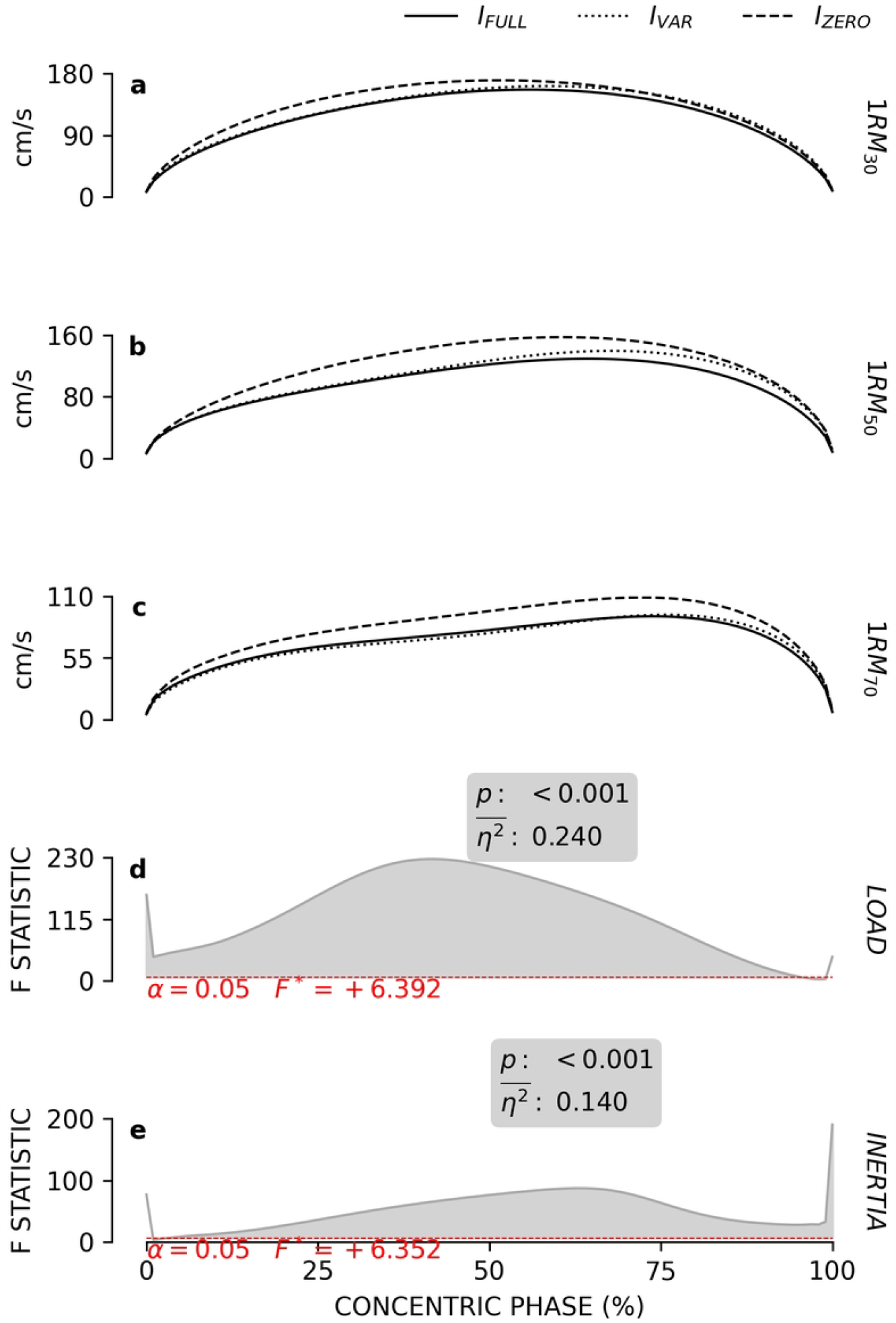
*VELOCITY* patterns. The subfigures **a, b** and **c** respectively report the mean velocity patterns obtained at 30, 50 and 70% of the *1RM* for each inertial profile. The grey lines in subfigures **d** and **e** show the statistical maps for the *LOAD* and *INERTIA* effects. The red dashes indicate the critical F value corresponding to *α*. The grey areas highlight the significant regions of the map and the boxes report the calculated *p* values and effect size 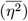.

*POWER* was influenced by *LOAD* on two separate *ROM* regions (Fig. 4). At *ROM* _15%:62%_, *1RM*_*70*_ resulted in lower values than *1RM*_*50*_ and *1RM*_*30*_. Conversely, over the *ROM* _93%:100%_ range, an opposite trend was found and *1RM*_*70*_ shown the highest *POWER* with respect to *1RM*_*30*_ and *1RM*_*50*_ (Table 1). *POWER* resulted affected by *INERTIA* at the end of the *ROM* (*ROM* _80%:100%_), where *I*_*VAR*_ achieving the highest values and *I*_*FULL*_ the lowest (Table 1).

**Fig 4.**
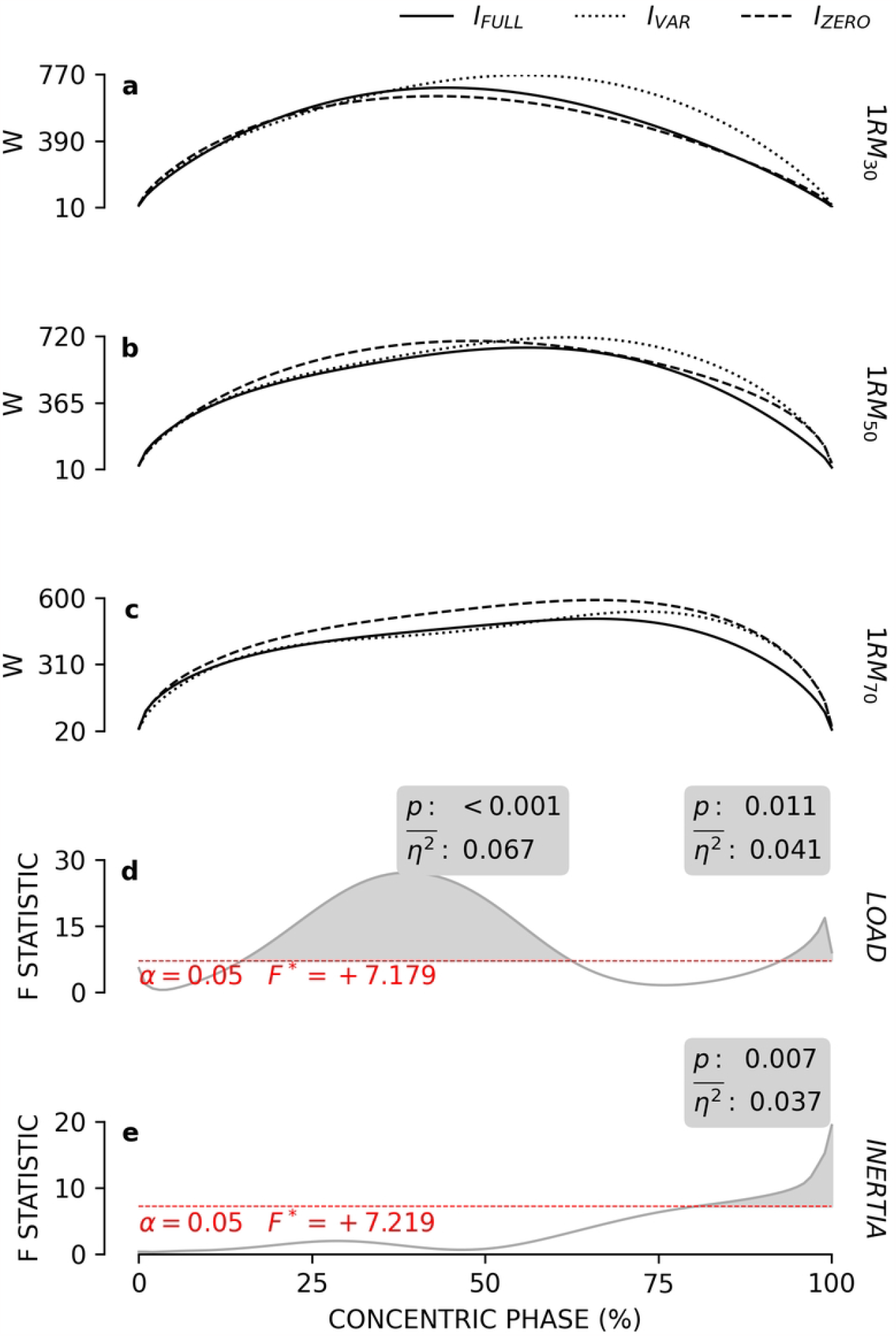
*POWER* patterns. The subfigures **a, b** and **c** respectively report the mean power patterns obtained at 30, 50 and 70% of the *1RM* for each inertial profile. The grey lines in subfigures **d** and **e** show the statistical maps for the *LOAD* and *INERTIA* effects. The red dashes indicate the critical F value corresponding to *α*. The grey areas highlight the significant regions of the map and the boxes report the calculated *p* values and effect size 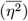.

*EMG*_*PM*_ amplitude resulted proportional to *LOAD* (Fig. 5) during the second half of the concentric phase (*ROM* _50%:97%_). Conversely, no effects were found for *INERTIA* in any part of the *ROM*.

**Fig 5.**
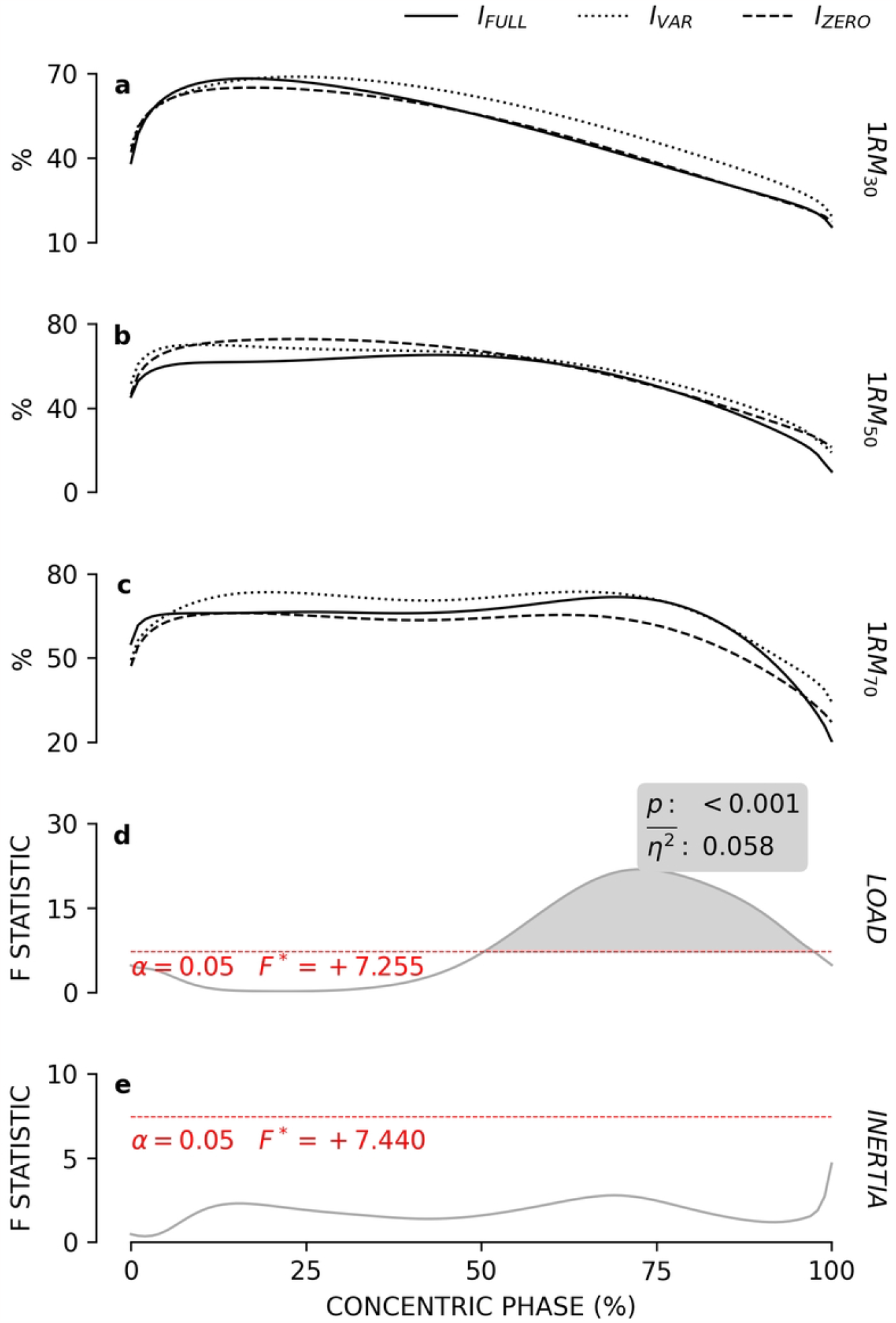
*EMG*_*PM*_ patterns. The subfigures **a, b** and **c** respectively report the mean *EMG* patterns of the *right pectoralis major* muscle obtained at 30, 50 and 70% of the *1RM* for each inertial profile. The grey lines in subfigures **d** and **e** show the statistical maps for the *LOAD* and *INERTIA* effects. The red dashes indicate the critical F value corresponding to *α*. The grey areas highlight the significant regions of the map and the boxes report the calculated *p* values and effect size 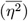.

*EMG*_*TB*_ (Fig. 6) activity was greater with higher loads over the large part of the *ROM* (*ROM* _14%:97%_). In addition, at the end of the concentric phase (*ROM* _98%:100%_), *I*_*FULL*_ lowered *EMG*_*TB*_ as compared to *I*_*ZERO*_ and *I*_*VAR*_ (Table 1).

**Fig 6.**
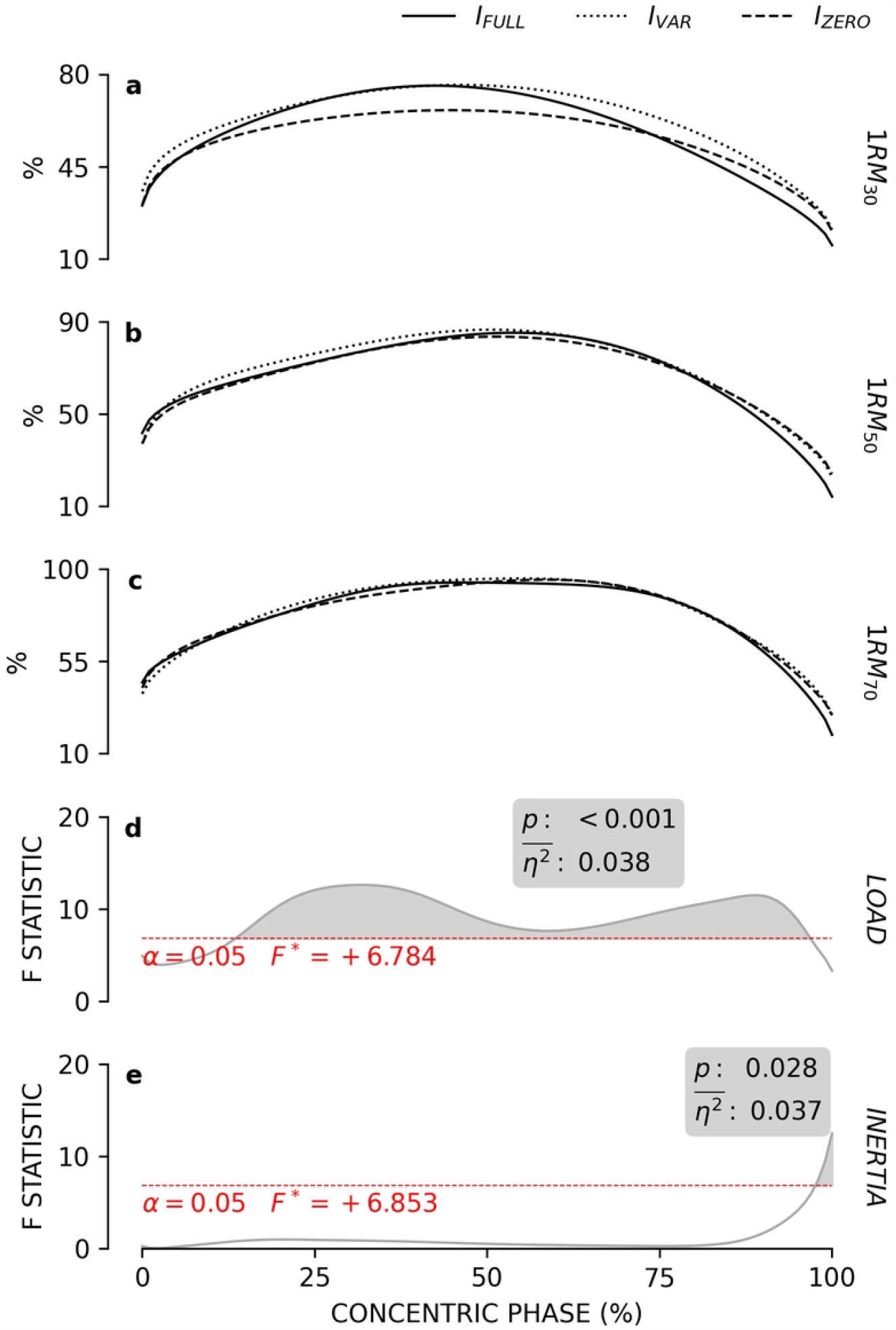
*EMG*_*TB*_ patterns. The subfigures **a, b** and **c** respectively report the mean *EMG* patterns of the *right triceps brachii* muscle obtained at 30, 50 and 70% of the *1RM* for each inertial profile. The grey lines in subfigures **d** and **e** show the statistical maps for the *LOAD* and *INERTIA* effects. The red dashes indicate the critical F value corresponding to *α*. The grey areas highlight the significant regions of the map and the boxes report the calculated *p* values and effect size 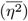.

*EMG*_*DA*_ was not influenced by *INERTIA* although *LOAD* shown significant effects over two regions of the *ROM* (Fig. 7). At *ROM* _0%:4%_, *1RM*_*30*_ resulted in lower *EMG* amplitude than *1RM*_*50*_ and *1RM*_*70*_ while at *ROM* _50%:94%_ the *EMG*_*DA*_ activity was proportional to *LOAD* (Table 1).

**Fig 7.**
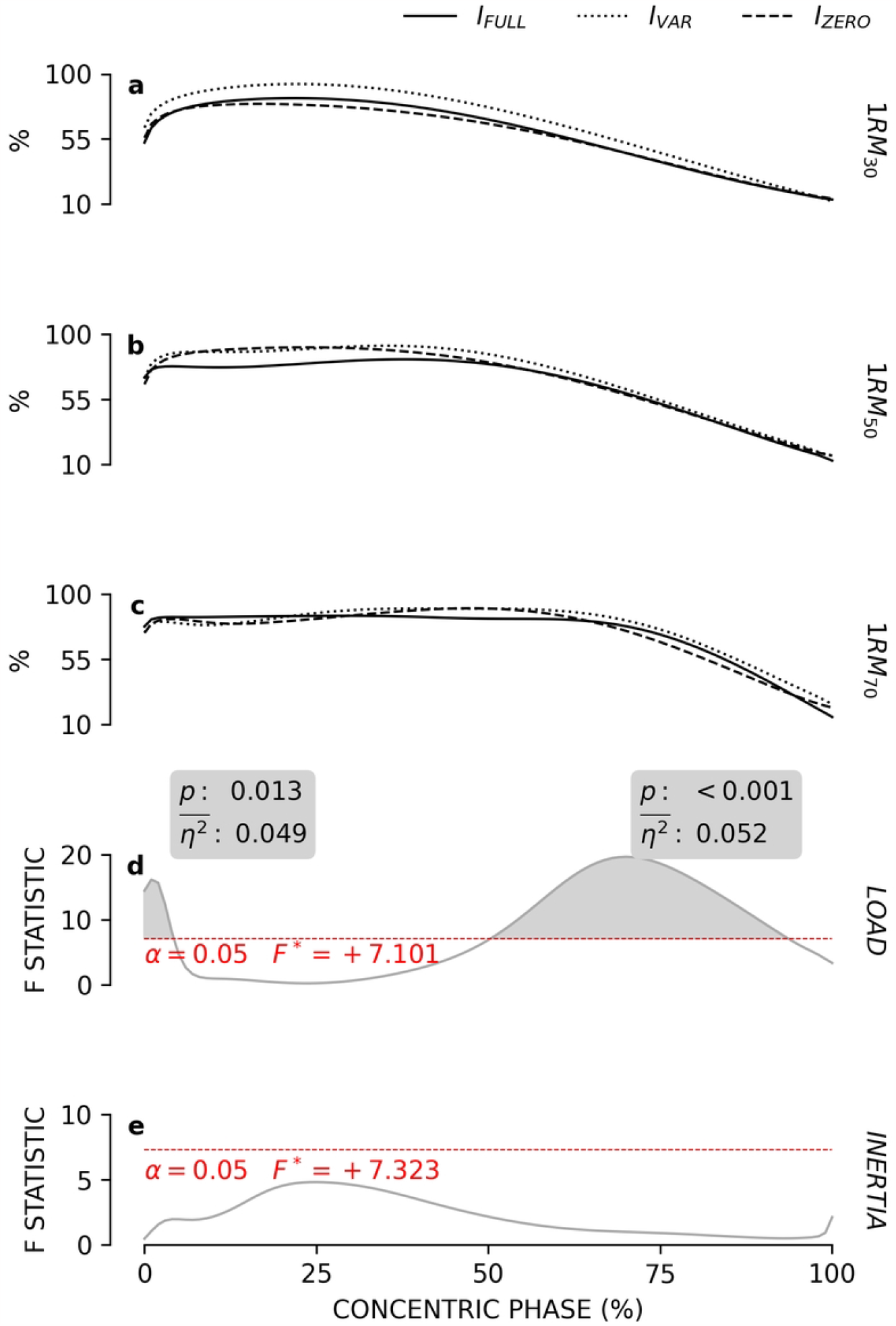
*EMG*_*DA*_ patterns. The subfigures **a, b** and **c** respectively report the mean *EMG* patterns of the *right anterior deltoid* muscle obtained at 30, 50 and 70% of the *1RM* for each inertial profile. The grey lines in subfigures **d** and **e** show the statistical maps for the *LOAD* and *INERTIA* effects. The red dashes indicate the critical F value corresponding to *α*. The grey areas highlight the significant regions of the map and the boxes report the calculated *p* values and effect size 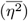.

The ^*<*xxx^ notation denotes significant lower values (*p <* 0.05) against the annotated condition.

## Discussion

This study examined the effects induced by different inertial resistance profiles on *FORCE, POWER, VELOCITY* and the amplitude of the *EMG*_*PM*_, *EMG*_*TB*_ and *EMG*_*DA*_ muscles during a chest press exercise performed at different intensity levels.

In line with other studies [27–29], higher intensity elevated *FORCE*. This effect was relevant across the whole *ROM* and acted independently to the employed inertial profile. As predicted by theory [14], *FORCE* was enhanced by the presence of inertial resistance at beginning of the *ROM*, and reduced at its end. As a result, the lack of inertial resistance (as per the *I*_*ZERO*_ condition) limited *FORCE*, although it provided a flatter profile across the *ROM*. To this extent, *I*_*VAR*_, who kept the inertial contribution of *I*_*FULL*_ at the beginning of the *ROM* and then progressively approached the *I*_*ZERO*_ behaviour toward the end of the concentric phase, retained elevated *FORCE* for longer portions of the *ROM* and remarkably overcame *I*_*ZERO*_ and *I*_*FULL*_ over the mid-range (Fig. 2).

In agreement with previous findings [12], the inclusion of inertial resistance reduced *VELOCITY* throughout the whole *ROM* (Fig. 3). The same findings supported the use of *I*_*ZERO*_ to achieve high *POWER* output. However, the results of this study suggests that inertial resistance is important for the maintenance of high *POWER* along the *ROM*. Notably, this study revealed that the *POWER* output was not affected by *INERTIA* over the first 80% of the *ROM*. Afterwards, *POWER* resulted progressively lower between *I*_*VAR*_, *I*_*ZERO*_ and *I*_*FULL*_ respectively (Fig. 4). This behaviour could be the result of the combination of effects already discussed about *FORCE* and *VELOCITY*. When *I*_*FULL*_ was employed, the relatively low *VELOCITY* was compensated by the higher *FORCE* levels. However, the reduction in *FORCE* induced by the facilitating effect of the inertial resistance at the end of the concentric phase had the effect of limiting the *POWER* output with respect to the other two inertial conditions. Following the same principle, the low *FORCE* of *I*_*ZERO*_ was compensated by the highest *VELOCITY* among the three tested inertial conditions. This allowed the production of high *POWER* levels throughout the major part of the *ROM*. Nevertheless, the higher *POWER* achieved over the last 20% of the *ROM* by *I*_*VAR*_ suggests also that the limited amount of *FORCE* produced with *I*_*ZERO*_ might have limited the *POWER* output over the later portion of the *ROM*. To this extent, while *I*_*ZERO*_ resulted the best choice for eliciting high *VELOCITY*, the possibility of varying the amount of inertial resistance throughout the *ROM* seemed to combine the positive effects of *I*_*FULL*_ on *FORCE* with those provided by *I*_*ZERO*_ on *POWER*.

Previous findings [11, 12] suggest that on the bench press exercise, in general the higher the *LOAD*, the higher the *EMG* amplitude produced by the prime movers. However, this rather simplistic rule is affected by other factors like execution speed and the type of resistance being employed. Indeed, the results of this study shown that higher *LOAD* did not change the *EMG*_*PM*_ amplitude over the first half of the *ROM*, but it required the *pectoralis major* muscle to be recruited for longer time (Fig. 5). As a result, elevated *EMG*_*PM*_ was found at higher *LOAD* only over the second half of the *ROM*. A very similar behaviour was found on the *anterior deltoid* muscle (Fig. 7) but not on the *triceps brachii*, which appeared more active as the load was increased over the major part of the *ROM* (Fig. 6). These results are well aligned to previous findings [30] indicating that, during the bench press concentric phase, the elbow extensors (i.e. the *triceps brachii* muscles) assist the action of the shoulder transversal flexors (e.g. the *pectoralis major* and *anterior deltoid* muscles) in driving away the barbell from the chest. The participants to this study were required to maximally accelerate the handles regardless to the load or the inertial resistance profile. This translates into the requirement of maximally accelerating the handles of the exercise apparatus, which in turn was designed to move along the shoulders transversal plane.

Accordingly, the *pectoralis major* and *anterior deltoid* activity resulted nearly maximal as the handles were accelerated independently to the load or inertial condition, while the contribution of *triceps brachii* increased proportionally to the moved resistance to support the other muscles through a more powerful elbow extension. This would explains also the lower *EMG*_*TB*_ amplitude found under the *I*_*FULL*_ condition at the very end of the *ROM*. As already discussed, when the movement starts to decelerate, the inertial resistance reduces the total amount of *FORCE* required. Accordingly, the lower *FORCE* requirement would have translated to a reduced effort imposed to the *triceps brachii* muscles and thus to the *EMG*_*TB*_ amplitude.

As a final remark, the possibility of varying the amount of inertia along the *ROM* is a novel opportunity allowed by the use of resistance exercise apparatuses driven by programmable electromechanical motors. In this study three simple inertial resistance profiles have been generated and compared, however, this novel technology might allow the design of more precise exercise routines and would require additional studies to better understand their potential.

## Conclusion

This study examined the effects of distinct inertial resistance profiles (*I*_*FULL*_, *I*_*ZERO*_ and *I*_*VAR*_) on *FORCE, POWER, VELOCITY* and on the *EMG*_*PM*_, *EMG*_*TB*_ and *EMG*_*DA*_ amplitude during a chest press exercise performed at different intensity levels. Compared to *I*_*FULL*_, *I*_*ZERO*_ allowed faster movements accompanied to lower *FORCE* outcomes throughout the whole *ROM*. Conversely *I*_*VAR*_, despite being slower than *I*_*ZERO*_, generated more *FORCE* and *POWER. POWER* was higher with lower loads in the mid-range of the *ROM*, while it was proportional to the applied resistance during the last part of the concentric phase. Higher loads elevated *EMG*_*PM*_ and *EMG*_*DA*_ mainly during the second half of the concentric phase, while *EMG*_*TB*_ was greater with heavier loads at the *ROM*’s mid-range. *I*_*FULL*_ reduced the *EMG*_*TB*_ amplitude at the end of the concentric phase, while its effect on *EMG*_*PM*_ and *EMG*_*DA*_ was negligible. As result, *I*_*VAR*_ overcame both *I*_*FULL*_ and *I*_*ZERO*_ in terms of *FORCE* and *POWER* output, which support the effectiveness of variable inertial profiles into resistance exercise programs. Ultimately, this study suggests that *INERTIA* is an independent factor of the *FORCE, VELOCITY* and *POWER* production. Its modulation should therefore be considered to adjust the effects induced by an exercise toward the achievement of higher *FORCE, POWER* or *VELOCITY*. Hence, coaches and therapists are encouraged to account for the type of *INERTIA* as one of the parameters considered during the preparation of the exercise routine for their athletes or patients.

## S1 Appendix

## Acknowledgments

The authors thank Mr. James Vernau for his helpful critical review of the study. All authors collaborated to the development of the exercise apparatus but none of the authors or participants received direct financial support for this study. All authors conceived and designed the research. The first author conducted the experiments and wrote the manuscript. All authors read, reviewed and approved the manuscript. The data presented in this study are available upon request to the corresponding author.

## References

1. Frost DM, Cronin J, Newton RU. A biomechanical evaluation of resistance: fundamental concepts for training and sports performance. Sports medicine (Auckland, NZ). 2010;40:303–26. doi:10.2165/11319420-000000000-00000.

2. Biscarini A, Bonafoni S. Optimization of the biomechanical design of plate-loaded strength training machines: The free-weight lifting experience. Proceedings of the Institution of Mechanical Engineers, Part P: Journal of Sports Engineering and Technology. 2017;231. doi:10.1177/1754337115624076.

3. Lesinski M, Prieske O, Granacher U. Effects and dose–response relationships of resistance training on physical performance in youth athletes: a systematic review and meta-analysis. British Journal of Sports Medicine. 2016;50:781–795. doi:10.1136/bjsports-2015-095497.

4. Garber CE, Blissmer B, Deschenes MR, Franklin BA, Lamonte MJ, Lee IM, et al. Quantity and Quality of Exercise for Developing and Maintaining Cardiorespiratory, Musculoskeletal, and Neuromotor Fitness in Apparently Healthy Adults. Medicine and Science in Sports and Exercise. 2011;43:1334–1359. doi:10.1249/MSS.0b013e318213fefb.

5. Biscarini A, Benvenuti P, Botti FM, Brunetti A, Brunetti O, Pettorossi VE. Voluntary enhanced cocontraction of hamstring muscles during open kinetic chain leg extension exercise: Its potential unloading effect on the anterior cruciate ligament. American Journal of Sports Medicine. 2014;42:2103–2112. doi:10.1177/0363546514536137.

6. Biscarini A. Measurement of power in selectorized strength-training equipment. Journal of Applied Biomechanics. 2012;28:229–241. doi:10.1123/jab.28.3.229.

7. Robertson DGE, Caldwell GE, Hamill J, Kamen G, N Whittlesey S. Research Methods in Biomechanics. Second edi ed. Human Kinetics; 2014.

8. Ratamess NA, Alvar BA, Evetoch TK, Housh TJ, Kibler WB, Kraemer WJ, et al. American College of Sports Medicine position stand. Progression models in resistance training for healthy adults. Medicine and science in sports and exercise. 2009;41:687–708. doi:10.1249/MSS.0b013e3181915670.

9. Davies TB, Kuang K, Orr R, Halaki M, Hackett D. Effect of Movement Velocity During Resistance Training on Dynamic Muscular Strength: A Systematic Review and Meta-Analysis. Sports Medicine. 2017;47:1603–1617. doi:10.1007/s40279-017-0676-4.

10. Ferri Marini C, Shoaei V, Micheli L, Francia P, Grossi T, Maggio S, et al. Barbell load distribution and lifting velocity affect bench press exercise volume and perceived exertion. PLOS ONE. 2022;17:e0278909. doi:10.1371/journal.pone.0278909.

11. Sakamoto A, Sinclair PJ, Moritani T. Muscle activations under varying lifting speeds and intensities during bench press. European Journal of Applied Physiology. 2012;112:1015–1025. doi:10.1007/s00421-011-2059-0.

12. Frost DM, Cronin JB, Newton RU. A comparison of the kinematics, kinetics and muscle activity between pneumatic and free weight resistance. European Journal of Applied Physiology. 2008;104:937–956. doi:10.1007/s00421-008-0821-8.

13. Anderson CE, Sforzo GA, Sigg JA. The effects of combining elastic and free weight resistance on strength and power in athletes. Journal of Strength and Conditioning Research. 2008;22:567–574. doi:10.1519/JSC.0B013E3181634D1E.

14. Biscarini A, Contemori S. Variable inertia training: Optimization of explosive-power exercises with robotic-resistance strength machines. Proceedings of the Institution of Mechanical Engineers, Part P: Journal of Sports Engineering and Technology. 2018;232:140–149. doi:10.1177/1754337117718086.

15. Sarto F, Franchi MV, Rigon PA, Grigoletto D, Zoffoli L, Zanuso S, et al. Muscle activation during leg-press exercise with or without eccentric overload. European Journal of Applied Physiology. 2020;120:1651–1656. doi:10.1007/s00421-020-04394-6.

16. Barbero M, Merletti R, Rainoldi A. Atlas of Muscle Innervation Zones. Springer Milan; 2012.

17. Coratella G, Tornatore G, Longo S, Esposito F, Cè E. Specific prime movers’ excitation during free-weight bench press variations and chest press machine in competitive bodybuilders. European Journal of Sport Science. 2020;20:571–579. doi:10.1080/17461391.2019.1655101.

18. Brzycki M. Strength Testing—Predicting a One-Rep Max from Reps-to-Fatigue. Journal of Physical Education, Recreation and Dance. 1993;64:88–90. doi:10.1080/07303084.1993.10606684.

19. Jidovtseff B, Harris NK, Crielaard JM, Cronin JB. Using the load-velocity relationship for 1RM prediction. Journal of Strength and Conditioning Research. 2011;25:267–270. doi:10.1519/JSC.0b013e3181b62c5f.

20. DeLuca CJ, Gilmore LD, Kuznetsov M, Roy SH. Filtering the surface EMG signal : Movement artifact and baseline noise contamination. Journal of Biomechanics. 2010;43:1573–1579. doi:10.1016/j.jbiomech.2010.01.027.

21. Thomas SJ, Zeni JA, Winter DA. Winter’s Biomechanics and Motor Control of Human Movement. 5th ed. Wiley; 2023.

22. McCaw ST, Friday JJ. A Comparison of Muscle Activity Between a Free Weight and Machine Bench Press. The Journal of Strength and Conditioning Research. 1994;8.

23. Pataky TC, Vanrenterghem J, Robinson MA. Two-way ANOVA for scalar trajectories, with experimental evidence of non-phasic interactions. Journal of Biomechanics. 2015;48:186–189. doi:10.1016/J.JBIOMECH.2014.10.013.

24. Jones RH. Bayesian information criterion for longitudinal and clustered data. Statistics in medicine. 2011;30:3050–6. doi:10.1002/sim.4323.

25. Pataky TC, Vanrenterghem J, Robinson MA. Zero-vs. one-dimensional, parametric vs. non-parametric, and confidence interval vs. hypothesis testing procedures in one-dimensional biomechanical trajectory analysis. Journal of biomechanics. 2015;48:1277–85. doi:10.1016/j.jbiomech.2015.02.051.

26. Holm S. A Simple Sequentially Rejective Multiple Test Procedure. Scandinavian Journal of Statistics. 1979;6:65–70.

27. Pearson SN, Cronin JB, Hume PA, Slyfield D. Kinematics and kinetics of the bench-press and bench-pull exercises in a strength-trained sporting population. Sports Biomechanics. 2009;8:245–254. doi:10.1080/14763140903229484.

28. Schilling BK, Falvo MJ, Chiu LZF. Force-velocity, impulse-momentum relationships: Implications for efficacy of purposefully slow resistance training. Journal of Sports Science and Medicine. 2008;7:299–304.

29. Picerno P, Iannetta D, Comotto S, Donati M, Pecoraro F, Zok M, et al. 1RM prediction: a novel methodology based on the force–velocity and load–velocity relationships. European Journal of Applied Physiology. 2016;116:2035–2043. doi:10.1007/s00421-016-3457-0.

30. Biscarini A, Calandra A, Contemori S. Three-dimensional mechanical modeling of the barbell bench press exercise: Unveiling the biomechanical function of the triceps brachii. Proceedings of the Institution of Mechanical Engineers, Part P: Journal of Sports Engineering and Technology. 2020;234:245–256. doi:10.1177/1754337120917831.

